# *In-vitro* and *in-vivo* characterization of CRANAD-2 for multi-spectral optoacoustic tomography and fluorescence imaging of amyloid-beta deposits in Alzheimer mice

**DOI:** 10.1101/2020.10.27.353862

**Authors:** Ruiqing Ni, Alessia Villois, Xose Luis Dean-Ben, Zhenyue Chen, Markus Vaas, Stavros Stavrakis, Gloria Shi, Andrew deMello, Chongzhao Ran, Daniel Razansky, Paolo Arosio, Jan Klohs

## Abstract

The abnormal deposition of fibrillar beta-amyloid (Aβ) deposits in the brain is one of the major histopathological hallmarks of Alzheimer’s disease (AD). Here we characterize curcumin-derivative CRANAD-2 for multi-spectral optoacoustic tomography (MSOT) and fluorescence imaging of brain Aβ deposits in the arcAβ mouse model of AD cerebral amyloidosis. CRANAD-2 shows a specific and quantitative detection of Aβ fibrils *in vitro,* even in complex mixtures, and it is capable to distinguish between monomeric and fibrillar forms of Aβ. *In vivo* epifluorescence and MSOT after intravenous CRANAD-2 administration demonstrated higher retention in arcAβ compared to non-transgenic littermate mice. Immunohistochemistry showed co-localization of CRANAD-2 and Aβ deposits in arcAβ mouse brain sections, thus verifying the specificity of the probe. In conclusion, we demonstrate suitability of CRANAD-2 for fluorescence- and MSOT-based detection of Aβ deposits in animal models of AD pathology, which facilitates mechanistic studies and the monitoring of putative treatments targeting Aβ deposits.

## 1. Introduction

The abnormal acculumation and the spread of amyloid-beta (Aβ) deposits play a central role in the pathogenesis of Alzheimer’s disease (AD) and leads to downstream pathophysiological events [1, 2]. Positron emission tomography (PET) imaging of aberrant Aβ deposits has been established as a diagnostic pathological biomarker for AD under clinical setting and been included in the new diagnostic criteria [3]. Three amyloid imaging probes have been approved for clinical usage including ^18^F-florbetapir [4], ^18^F-fl orb etab en [5] and ^18^F-flumetamol [6]. Higher PET imaging of cortical Aβ loads were reported in the brains from patients with AD and mild cognitive impairment compared to healthy controls [7, 8]. While PET imaging in small rodents has been used for studying Aβ-related disease mechanisms and for developing therapeutic strategies [9–12], its resolution (1 mm) relative to the mouse brain dimension (10 mm × 8 mm) and the low availability of radiotracer and imaging facilities restrict its wide usage.

Optical imaging techniques such as near-infrared fluorescence (NIRF) imaging and multi-spectral optoacoustic tomography (MSOT) imaging have emerged, enabling the *in vivo* imaging of physiological, metabolic and molecular function [13–16]. This is mainly due to the fact that optical techniques have a high detection sensitivity, are cost-effective and use nonionizing radiation. Progress is made not only in the development of optical instrumentation and reconstruction algorithms, but also in the design and synthesis of novel Aβ imaging probes [17]. For example, several probes have been reported for near-infrared fluorescence imaging including NIAD-4 [18], AOI987 [19], BODIPY [20], THK-265 [21], DANIR [22–25], curcumin-derivatives CRANAD series [26, 27], luminescent conjugated oligothiophenes [28, 29] and DBA-SLOH [30]. NIRF imaging of Aβ deposits has been shown using a variety of AD mouse models [19, 23–27, 30, 31]. Despite its advantages, the diffuse nature of light propagation in highly scattering tissue prevents the accurate determination of probe distribution and concentration from NIRF measurements. As an inherent hybrid method, MSOT imaging capitalizes on absorption of light as a source of contrast, while emitted non-radioactive decay broadband ultrasound is used for image formation, thus combining the high sensitivity of optical imaging with the high spatial resolution of ultrasound imaging. Recently, volumetric multispectral optoacoustic tomography (vMSOT) has been introduced with added multiplexing and real-time 3D imaging capabilities, thus enabling a wide range of new biomedical applications [32–37]. Previously, intravital optoacoustic imaging of Aβ deposits with intrathecal injection of Congo Red [27] and croconium-derivative [38] in amyloidosis mouse models have been reported.

In the present study, we show that CRANAD-2, previously used for NIRF imaging [26, 27], is suitable for MSOT imaging and describe its utility for *in vivo* whole brain mapping of Aβ deposits in arcAβ mouse model of cerebral amyloidosis [39]. Capitalizing on the fluorescent properties we characterized binding of the probe to Aβ fibrils *in vitro* using a fluorescence binding assays, *in situ* with tissue sections and immunohistochemistry and by cross-validation between MSOT and hybrid fluorescence imaging [40, 41]. Our results demonstrate suitability of CRANAD-2 for NIRF and MSOT imaging of Aβ deposits.

## 2. Materials and methods

### 2.1 Animal model

Seven transgenic arcAβ mice overexpressing the human APP695 transgene containing the Swedish (K670N/M671L) and Arctic (E693G) mutations under the control of prion protein promoter and five age-matched non-transgenic littermates of both sexes were used (18-24 months-of-age) [39]. ArcAβ mice are characterized by a pronounced amyloid deposition, cerebral amyloid angiopathy and vascular dysfunction [36, 42–46], Animals were housed in ventilated cages inside a temperature-controlled room, under a 12-hour dark/light cycle. Pelleted food (3437PXL15, CARGILL) and water were provided *ad-libitum.* Paper tissue and red Tecniplast mouse house® (Tecniplast, Milan, Italy) shelters were placed in cages as environmental enrichments. All experiments were performed in accordance with the Swiss Federal Act on Animal Protection and were approved by the Cantonal Veterinary Office Zurich (permit number: ZH082/18).

### 2.2 *In vitro* binding between amyloid probes and recombinant Aβ_1-42_ fibrils measured by spectrofluorometer

(T-4)-[(1E,6E)-1,7-Bis[4-(dimethylamino)phenyl]-1,6-heptadiene-3’5-dionato-kO3’kO_5_] difluoroboron (CRANAD-2) [26] was purchased from Sigma-Aldrich AG, Switzerland. Recombinant Aβ1.42 monomers were expressed and produced by *E.coli* as described previously [47]. Aβ1.42 fibrils were formed by incubating a solution of 2 μM Aβ1.42 in phosphate buffer (20 mM sodium phosphate, 0.2 mM ethylenediaminetetraacetic acid, pH 8.0). The aggregation process was monitored by a quantitative fluorescence assay based on the Thioflavin T (ThT) dye [48]. CRANAD-2 stock was dissolved in dimethyl sulfoxide (DMSO, Sigma-Aldrich AG). The binding *in vitro* between CRANAD-2 (100 nM) or ThT (200 μM) with Aβ_1-42_ fibrils (0 - 200 nM) was measured on a spectrofluorometer (Fluoromax-4, HORIBA Jobin Yvon Technologies, Japan) by recording the emission spectra from 660 nm - 800 nm after excitation at 640 nm or 450 nm - 650 nm respectively. To investigate whether CRANAD-2 binds selectively Aβ_1-42_ fibrils, we assessed the binding of CRANAD-2 (100 nM) and ThT (200 μM) to other targets that are rich in β-sheet structures including bovine serum albumin, monoclonal antibody, and amyloids formed by lysozyme, α-synuclein and insulin, all at 200 nM. Moreover, we assessed whether CRANAD-2 can distinguish between different conformations of Aβ by measuring binding also to Aβμ_42_ monomers. Finally, the specificity of CRANAD-2 for Aβ_ļ-42_ fibrils was further evaluated by measuring the binding between the probe and Aβ_1-42_ fibrils (0-200 nM) spiked in a cell lysate (with a total protein concentration of 7 μM).

### 2.3 NIRF imaging

NIRF imaging were performed with the Maestro 500 multispectral imaging system (Cambridge Research & Instruments Inc.). For phantom measurements, tubes (3 mm diameter, 4 to 5 cm lengths) were filled with either 200 μl Aβ fibrils (4 μM), or CRANAD-2 (30 μM), or CRANAD-2(30 μM) + Aβ (4 μM) samples.

### 2.4 MSOT

Cross-sectional MSOT imaging was performed with a commercial inVision 128 small animal scanner (iThera Medical, Germany) as described [49]. Briefly, a tunable (680-980 nm) optical parametric oscillator pumped by an Nd:YAG laser provides 9 ns excitation pulses at a framerate of 10 Hz with a wavelength tuning speed of 10 ms and a peak pulse energy of 100 mJ at 730 nm. Ten arms, each containing an optical fiber bundle, provide even illumination of a ring-shaped light strip with a width of approx. 8 mm. For ultrasound detection, 128 cylindrically focused ultrasound transducers with a center frequency of 5 MHz (60 % bandwidth), organized in a concave array of 270 degree angular coverage and a curvature radius of 4 cm, were used. Phantom and *in vivo* MSOT images were acquired at 10 wavelengths, i.e. 680, 685, 690, 695, 700, 715, 730, 760, 800 and 850 nm, coronal section, averages = 10, field-of-view = 20 × 20 mm, resolution = 100 μm × 100 μm, step = 0.3 mm moving along the axial direction.

For phantom measurements, tubes (3 mm diameter, 4 to 5 cm lengths) were filled with either 200 μl Aβ fibrils (4 μM), or CRANAD-2 (30 μM), or CRANAD-2 (30 μM) + Aβ (4 μM) samples. Tubes were placed in a cylindrical phantom (2 cm diameter, 2 % agarose mixed with 5 % intralipid (Sigma-Aldrich AG, Switzerland). The phantom holder was placed in an imaging chamber filled with water and kept at 28°C.

For *in vivo* MSOT, five arcAβ mice and four non-transgenic littermates were imaged with MSOT *in vivo.* Mice underwent also magnetic resonance imaging (MRI) for co-registration with MSOT data and to facilitate volume-of-interest analysis. Mice were anesthetized with an initial dose of 4 % isoflurane (Abbott, Cham, Switzerland) in oxygen/air (200/800 mL/min) mixture and were maintained at 1.5 % isoflurane in oxygen/air (100/400 mL/min). The fur over the head was epilated. The mouse was placed in a mouse holder in prone position. The holder was inserted in an imaging chamber filled with water to keep body temperature within 36.5 ± 0.5 °C. Mice were injected intravenously with CRANAD-2 (2.0 mg/kg, 15 % DMSO + 70 % PBS pH 7.4 + 15 % Kolliphor EL, Sigma-Aldrich AG) through the tail vein. Datasets were recorded before, 20, 40, 60, 90, and 120 minutes after the injection.

MSOT images were reconstructed using a model-based algorithm, size 20 mm, resolution 100 μm, and filter range from 50 kHz to 7 MHz. The model-based reconstruction incorporates a detailed model of detection geometry that allows for more quantifiable images.

### 2.5 Hybrid vMSOT-fluorescence imaging

A hybrid vMSOT-fluorescence imaging system [32, 50] was used to assess one arcAβ mouse. A short-pulsed (<10 ns) laser was used to provide an approximately uniform illumination profile on the mouse brain surface with optical fluence <20 mJ/cm^2^. The excited OA responses were collected with a custom-made spherical array (Imasonic SaS, Voray, France) of 512 ultrasound detection elements with 7 MHz central frequency and > 80 % bandwidth. A custom-made optical fiber bundle (Ceramoptec GmbH, Bonn, Germany) with 4 outputs was used for guiding the laser beam from multiple angles through the lateral apertures of the array. The detected signals were digitized at 40 megasamples per second with a custom-made data acquisition system (DAQ, Falkenstein Mikrosysteme GmbH, Taufkirchen, Germany) triggered with the Q-switch output of the laser. The pulse repetition frequency of the laser was set to 100 Hz and the laser wavelength tuned between 660 and 850 nm (10 nm step) on a per pulse basis. For the concurrent 2D epifluorescence imaging [41], beam from the pulsed OPO laser was similarly used for excitation. The generated fluorescence field was collected by a fiberscopic imaging fiber bundle comprised of 100,000 fibers and then projected onto an EMCCD camera (Andor Luca R, Oxford Instruments, UK). For hybrid imaging, the fiberscope was inserted into the central aperture of the spherical array detector. The system has an overall field-of-view of 15×15×15mm^3^ and resolution in the 110 μm range for both vMSOT and epifluorescence measurements [32, 51]. The optoacoustic signals were recorded before injection, 20, 40, 60, 90 and 120 min after injection of CRANAD-2; fluorescence signals were recorded before injection, 22, 42, 62, 92 and 120 minutes after injection of CRANAD-2. To examine the influence of scalp on the absorbance intensity, the scalp of the mouse was then removed. The mouse head was imaged afterwards using the same setting.

For the hybrid vMSOT-fluorescence measurements, images were rendered in real-time during the acquisition via fast back-projection-based image reconstruction implemented on a graphics processing unit [52]. A 3D model-based iterative algorithm was used off-line for more accurate reconstruction [53]. Prior to reconstruction, the collected signals were band-pass filtered with cut-off frequencies of 0.1 and 9 MHz. However, acoustic distortions associated to speed of sound heterogeneities, acoustic scattering and attenuation as well as the response of the ultrasound sensing elements of the array are known to additionally play a role in the quality of the images rendered [54].

### 2.6 MRI

MRI scans were performed on a 7/16 small animal MR Pharmascan (Bruker Biospin GmbH, Ettlingen, Germany) equipped with an actively shielded gradient capable of switching 760 mT/m with an 80-μs rise time and operated by a ParaVision 6.0 software platform (Bruker Biospin GmbH, Ettlingen, Germany). A circular polarized volume resonator was used for signal transmission, and an actively decoupled mouse brain quadrature surface coil with integrated combiner and preamplifier was used for signal receiving. Mice were anesthetized with an initial dose of 4% isoflurane (Abbott, Cham, Switzerland) in oxygen/air (200/800 mL/min) mixture and were maintained at 1.5% isoflurane in oxygen/air (100-400 mL/min). Mice were next placed in prone position on a water-heated support to keep body temperature within 36.5°C ± 0.5°C, monitored with a rectal temperature probe. *In vivo* T_2_-weighted MR images of mouse brain/head were obtained using a 2D spin echo sequence (Turbo rapid acquisition with refocused echoes) with imaging parameters: RARE factor 8, echo time 36 ms, repetition time 2628 ms, 6 averages, slice thickness 0.7 mm, no slice gap, field of view 20 × 20 mm2, matrix 512 × 512, giving an in-plane spatial resolution 39 × 39 μm^2^, out-of-plane resolution 0.7 mm (slice thickness), within a scan time 12 min 36 s.

### 2.7 Co-registration with MRI and volume-of-interest analysis of MSOT data

Registration between MSOT data and mouse brain MR images can provide a better anatomical reference for regional analysis. The cross-sectional MSOT images were co-registered with T_2_ - weighted MRI data as described [55]. For dynamic data, time course of regional absorbance (a.u.) at 680 nm were plotted and the area-under-curves were calculated. The retention of probe was calculated by averaging values from 60-120 min post-injection.

### 2.8 Immunohistochemistry and confocal imaging

For immunohistochemistry and confocal microscopy, mice were perfused with 1 × PBS (pH 7.4), under ketamine/xylazine/acepromazine maleate anesthesia (75/10/2 mg/kg body weight, bolus injection) and decapitated. Brains were removed from the skull afterwards, fixed in 4 % paraformaldehyde in 1 × PBS (pH 7.4). Brain tissue were cut horizontally at 5 μm and stained with CRANAD-2, anti-Aβ_1-16_ antibody 6E10 (Signet Lab, SIG-39320, 1:5000), fibrillar conformation anti-amyloid antibody OC (Merck, AB2286, 1:200), Donkey-anti-Rat Alexa 488 (Jackson, AB-2340686, 1:400), Goat-anti-Rabbit Alexa488 (Invitrogen, A11034, 1:200) and counter-stained using 4’,6-diamidino-2-phenylindole (DAPI) for nuclei (Sigma, D9542-10MG, 1:1000). Confocal images of the cortex of non-transgenic littermates and arcAβ mice were obtained using a Leica SP8 confocal microscope (Leica Microsystems GmbH, Germany) at ScopeM ETH Zurich Hönggerberg core facility. Sequential images were obtained by using 458 nm, 640 nm lines of the laser respectively. Identical settings were used and images were obtained for the non-transgenic littermates and arcAβ mice at ×20 and ×60 magnification, resolution with Z stack and maximum intensity projection.

### 2.9 Statistics

Unpaired two-tail *student t* test with Welch’s correction was used (Graphpad Prism 8.2, Graphpad Software Inc, USA) for comparing values between arcAβ mice and non-transgenic littermates. All data are present as mean ± standard deviation. Significance was set at * *p* < 0.05.

## 3 Results

### 3.1. *In vitro* characterization of probes on recombinant Aβ_1-42_ fibrils

We performed a fluorescence binding assay to assess the specificity of CRANAD-2 to Aβ fibrils (**Fig. 1**) and compare with the Thioflavin T (ThT) dye, a small molecule that gives strong fluorescence upon binding to amyloids and is currently the most common probe for *in vitro* assay of amyloid formation [56]. As expected, ThT binding to aggregated Aβ_1-42_ fibrils (0-100 nM), resulting in a dose-dependent increase in fluorescence intensities at X nm (**Fig. 1a**). In comparison, upon binding to aggregated Aβ_1-42_ fibrils CRANAD-2 showed an increase in fluorescence intensity at 681 nm (**Fig. 1d**). Linear relations between fluorescence intensity and concentration of aggregated Aβ_1-42_ fibrils (0-100 nM) were observed with CRANAD-2 (100 nM, **Figs. 1e-f**, *r* = 0.993) and ThT (200 μm, **Figs. 1b-c**, *r* = 0.955). In the presence of 200 nM Aβ_1-42_ fibrils the intensity is approximately 25-fold higher for CRANAD-2 compared to ThT (2.01×10^3^ *vs* 4.99×10^4^ counts per seconds).

**Fig 1.**
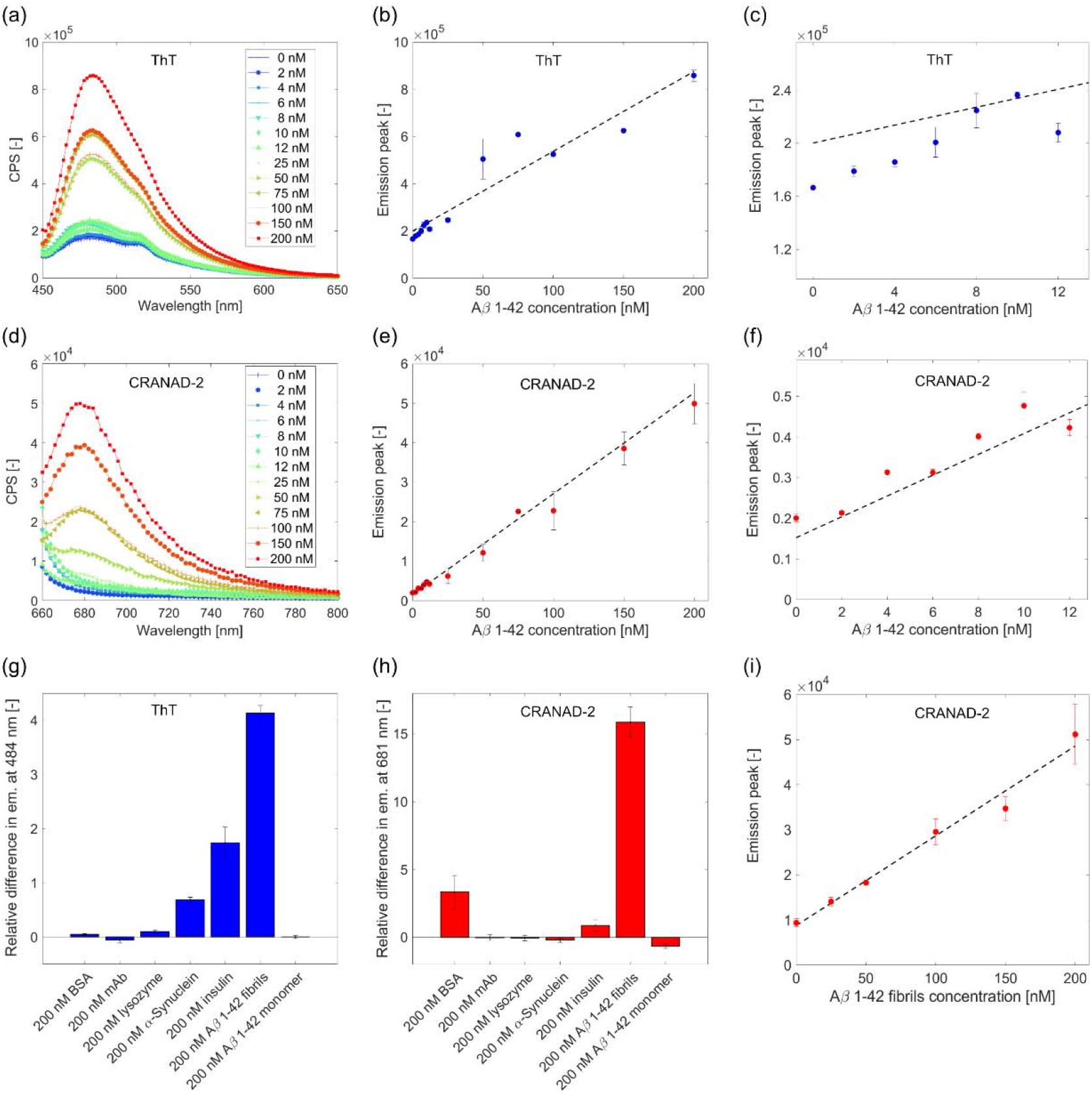
*In vitro* characterization of CRANAD-2 binding on recombinant Aβ_1-42_fibrils. (**a-f**) Fluorescence intensity (counts per seconds, CPS) of recombinant Aβι_-42_ fibrils (0-200 nM) in the presence of ThT (**a**) and CRANAD-2 (**d**); Linear relation between emission peak of ThT (484 nm, **b** and zoom-in **c**) and CRANAD-2 (681 nm, **e** and zoom-in **f**) with Aβ_1-42_ fibrils concentration; Comparison of ThT (**g**) and CRANAD-2 (**h**) binding to different proteins. BSA: bovine serum albumin; mAb: monoclonal antibody; (**i**) Linear relation of the fluorescence intensity of CRANAD-2 as function of Aβ_1-42_ fibrils spike in a cell lysate;

In a second set of experiments, we assessed the binding of CRANAD-2 and ThT to other proteins, including monomeric proteins rich in β–sheet structures, amyloid fibrils composed of other proteins and monomeric Aβ_1-42_ peptide. ThT showed specific binding to Aβ_1-42_ fibrils, a lower binding to α-synuclein, and insulin (**Fig. 1g**).

CRANAD-2 showed higher intensity signal upon binding to Aβ_1-42_ fibrils compared to fibrils composed of α-synuclein, insulin, lysosome, and to the Aβ_1-42_ monomer, while a non-negligible signal was observed in the presence of bovine serum albumin (**Fig. 1h**). Also ThT showed a higher signal upon binding to Aβ_1-42_ fibrils but compared to CRANAD-2 the intensity upon binding to fibrils of α-synuclein, and insulin was higher (**Fig. 1g**). Therefore, with respect to ThT, CRANAD-2 shows more specificity to Aβ_1-42_ fibrils and less generic binding to β-Sheet structures.

We further investigated the specificity of CRANAD-2 (100 nM) for Aβ_1-42_ fibrils (0-200 nM) in a complex sample (cell lysate with estimated total protein concentration of 7 μM). Notably, a linear relationship between Aβ_1-42_ fibrils and CRANAD-2 fluorescence intensity was observed even when fibrils were spiked in a cell lysate with much higher concentration of other proteins (**Fig. 1i**).

Overall, these results demonstrate that CRANAD-2 can bind Aβ_1-42_ fibrils with high affinity, specificity and selectivity. Importantly, the probe can distinguish between monomeric and fibrillar forms of the peptide.

### 3.2. *In vitro* binding of CRANAD-2 to Aβ fibrils increases NIRF and OA signal

CRANAD-2 was incubated with Aβ fibrils and measured in an agar phantom with NIRF imaging and MSOT (**Fig. 1**). Analysis of NIRF images of the samples indicated a 50 % increase in fluorescence intensity when CRANAD-2 was co-incubated with Aβ fibrils (**Figs. 2a, b**). In comparison, analysis of the probe phantom measured with MSOT showed a 100 % increase in the OA signal (**Figs. 2a, c**).

**Fig 2.**
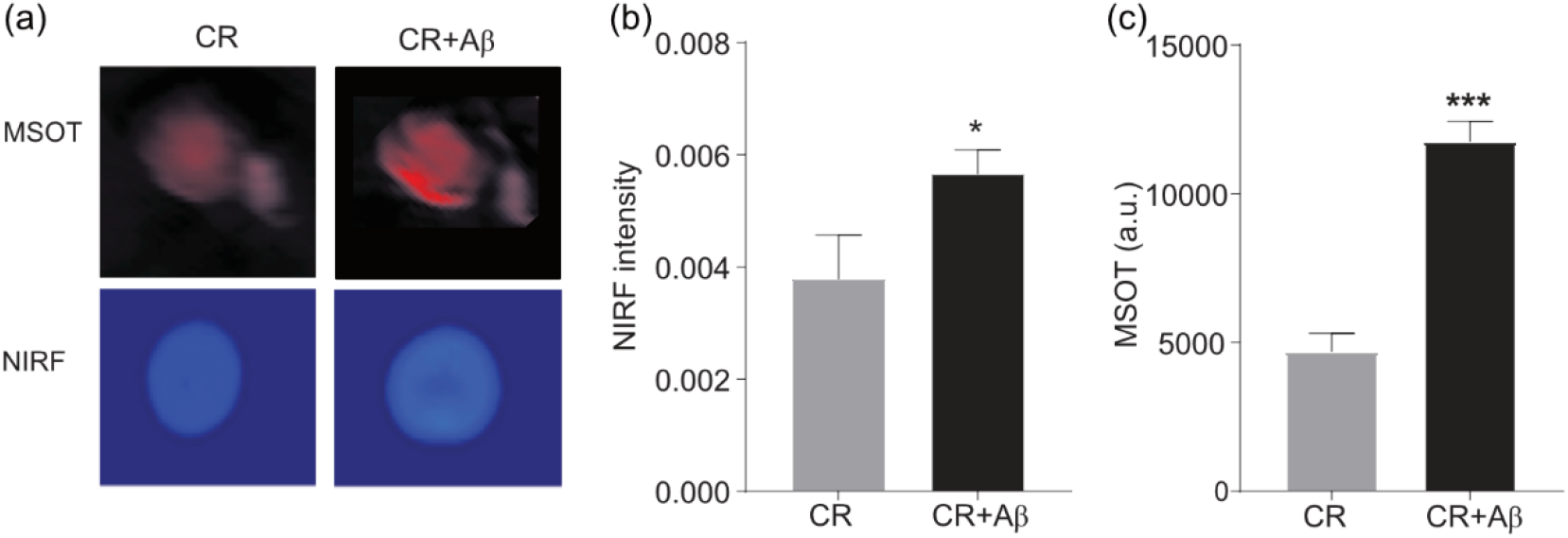
*In vitro* fluorescence and optoacoustic imaging of CRANAD-2 and Aβ fibrils. **(a)** Tubes containing CRANAD-2 (30 μM) and CRANAD-2 (30 μM) **+**Aβ (4 μM) inserts measured by using MSOT (in agar phantom) and NIRF imaging; (**b, c**) Quantification of NIRF and MSOT signal intensity. CR: CRANAD-2.

### 3.3. MSOT detects Aβ deposits *in vivo*

CRANAD-2 has been shown to cross the blood-brain-barrier and reach the brain parenchyma [26, 27]. We explored the ability of OA imaging to detect Aβ deposits *in vivo* in mouse brain. This, we set out to explore the ability of 3D whole brain Aβ mapping with MSOT. We monitored CRANAD-2 uptake in arcAβ mice and non-transgenic littermates for 120 min post-injection (**Fig. 3**). The delta images in relation to the pre-injection images shows high probe uptake in the brain of arcAβ mice, mainly in the cortex (**Fig. 3a**). Analysis of the dynamics of cortical CRANAD-2 uptake allowed to discriminate arcAβ mice from non-transgenic littermates (**Fig. 3**). The signal plateaued around 90 minutes, where higher AUC was observed in the cortical region of arcAβ mice compared to non-transgenic littermates (**Fig. 3c**).

**Fig. 3.**
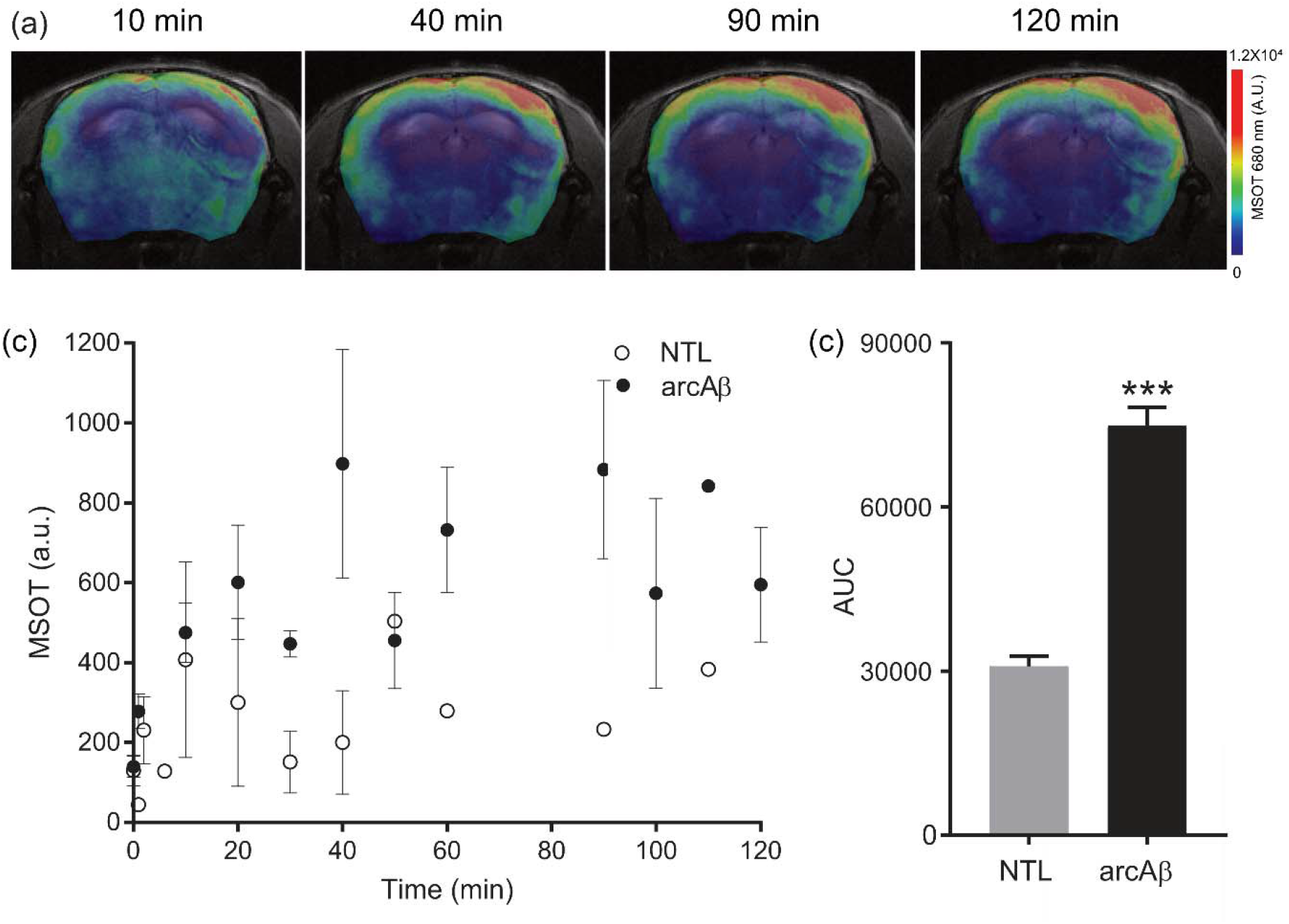
*In vivo* optoacoustic imaging of CRANAD-2 in mouse models. **(a)** Coronal view of optoacoustic images of arcAβ mice (overlaid with T_2_-weighted structural MRI) at 680 nm at 10, 40, 90 and 120 min after *i.v.* tail vein injection of CRANAD-2; (**c**) Time course of MOST intensity at 680 nm in the cortex of NTL and arcAβ mice over 120 minutes; (**d**) Quantification of area-under-curve (AUC) in NTL and arcAβ mice.

### 3.4. *In vivo* hybrid vMSOT and epifluorescence imaging in arcAβ mouse brain

We used a hybrid system to assess simultaneous vMSOT and epifluorescence imaging for CRANAD-2 distribution in arcAβ mouse brain. One arcAβ mouse was scanned *in vivo* using the hybrid system with intact scalp till 120 minutes after CRANAD-2 probe injection. A measurement was made without scalp at the end of data acquisition at 140 minutes (**Fig. 4**). We did not attain actual unmixing of CRANAD-2 (probably due to the similarity in absorbance spectrum with deoxyhemoglobin), and deoxy/oxyhemoglobin signal. The fluorescent CRANAD-2 spots are clearly visible with scalp (**Figs. 4a, b**) and without scalp (**Figs. 4a, d**). In comparison to the fluorescence signal, the degree of changes in MSOT signal (at single 660 nm wavelength) in the arcAβ mouse brain was moderate (**Figs. 4b, e**).

**Fig 4.**
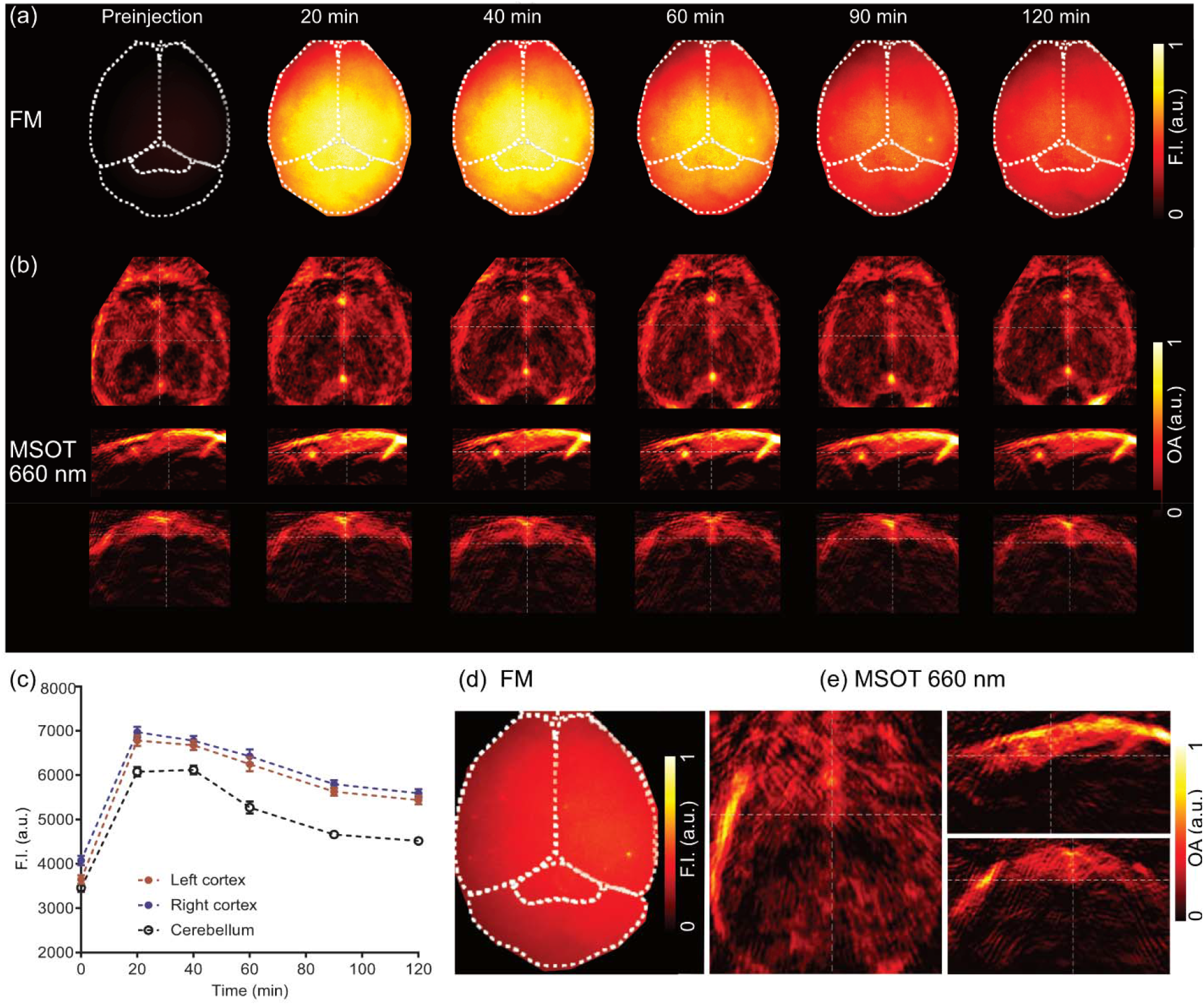
*In vivo* hybrid vMSOT and epifluorescence imaging of CRANAD-2 in an arcAβ mouse. (**a**) Epifluorescence images (horizontal); (**b**) corresponding MSOT images at 660 nm (horizontal, sagittal and coronal views) are shown respectively; at pre-injection, 20 to 120 min post-injection of CRANAD-2; (**c**) Fluorescence signal as a function of time in the right and left cortex and compared to cerebellum of arcAβ mouse; (**d**) Fluorescence (horizontal view) and (**e**) MSOT signal at 660 nm (horizontal, sagittal and coronal views) when the scalp was removed at 140 min post-injection.

### 3.5. *Ex vivo* staining on mouse brain sections

To validate the specificity of CRANDA-2 binding to Aβ deposits in mouse brain, horizontal brain tissue sections from arcAβ mice and non-transgenic littermates were stained with CRANAD-2 in addition to anti-Aβ antibodies 6E10, which binds any type of Aβ, and OC, which recognizes mature fibrils [57], and were nuclear counterstained with DAPI (**Fig. 5**). No signal was observed in the cortex from non-transgenic littermates (**Figs. 5a, f**). CRANAD-2 clearly costained with OC or 6E10 stained parenchymal and vessel wall-associated Aβ deposits in the arcAβ mouse brain (**Figs. 5b-e, g-n**). This suggests specific binding of CRANAD-2 to Aβ deposits. We also observed auto-fluorescence (blue) of Aβ plaques in the cortical region of arcAβ mouse brain tissue slice (**Fig. 5o**), similar to other reports [58].

**Figure 5.**
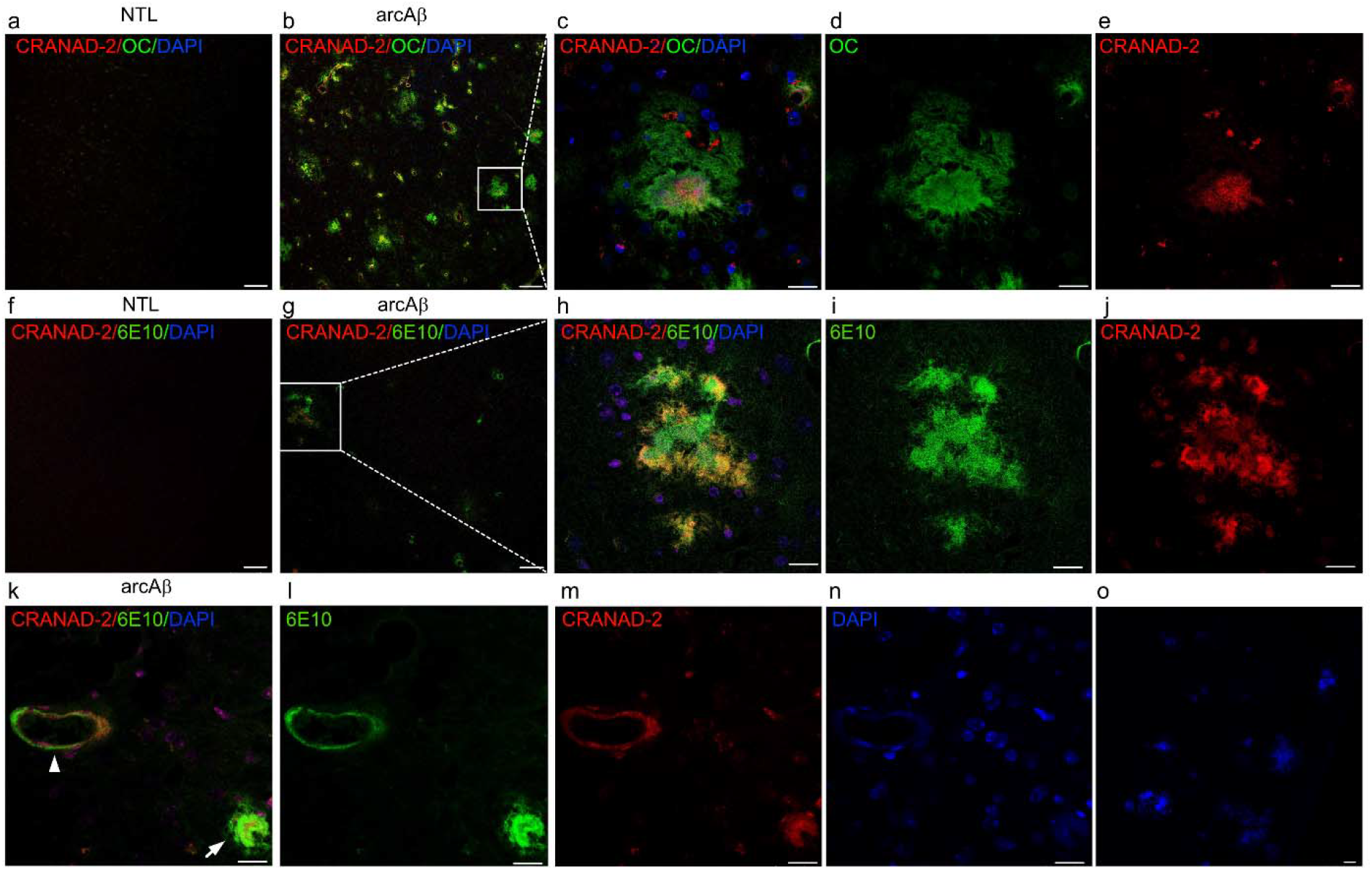
Co-staining of CRANAD-2 with Aβ deposition in non-transgenic littermate and arcAβ mouse brain tissue sections. Sections were stained with CRANAD-2 (red) and fibrillar conformation antibody OC (green) and anti-Aβ_1–16_ antibody 6E10 (green), and DAPI (blue) in the cortex of non-transgenic littermates (NTL) (**a, f**) and arcAβ mice (**b, g**); (**c-e, h-j**) zoom-in illustrating co-localization of the signal in (**b, g**); (**k-n**) Vessel associated cerebral amyloid angiopathy (arrowhead) and parenchymal plaque (arrow); (**o**) Autofluorescnce of Aβ plaques in the cortex of arcAβ mouse in the absence of stainings; Scale bars = 50 μm (**a, b, f, g**), 20 μm (**c-e, h-o**).

## 4 Discussion

Developing tools for non-invasive detection of Aβ deposits at high-resolution in animal models of AD is critical for investigating disease mechanism and translational development of Aβ-targeted therapies. Most probes for Aβ including NIAD-4 [18], AOI987 [19], BODIPY [20], THK-265 [21], DANIR [22–25], curcumin-derivatives CRANAD series [26, 27], luminescent conjugated oligothiophenes [28, 29] and DBA-SLOH [30] have been so far been designed for fluorescence imaging application. Nevertheless, Aβ binding fluorescent probes with suitable absorbance spectrum can be employed for OA imaging. We took advantage of the fluorescent property of CRANAD-2 to characterize the suitable of the probe to map cerebral Aβ deposits with NIRF and OA imaging.

CRANAD-2 has an affinity to Aβ aggregates of 38.7 nM [26], which is much higher than those of BODIPY, AOI-987 [19], similar to DANIR [25] and lower than NIAD-4 [18, 25]. The *in vitro* binding assay demonstrated a linear relationship between fluorescence intensity and concentration of aggregated Aβ_1-42_ fibrils for CRANAD-2. This properties has been reported for many Aβ binding probes such as AOI987, THK-265, DANIR and croconium dye [19, 21, 25, 38] and constitutes an advantage for the detection of fluorescence in NIRF imaging. However, when applying such probes for MSOT imaging, the increase in fluorescence quantum yield upon Aβ binding leads to a decrease in MSOT signal. Despite this feature, two previous studies in a transgenic mouse model of amyloidosis have reported Aβ imaging with other optoacoustic imaging systems using Congo Red [27], and croconium dye [38]. We were also able to clearly detect optoacoustic signals from CRANAD-2 bound to Aβ fibrils in the phantom and in arcAβ mice *in vivo.* Moreover, in the phantom the detected signal increased upon binding of CRANAD-2 to recombinant Aβ fibrils, which was higher for MSOT than NIRF imaging. In addition to an increase in fluorescence intensity, we observed a slight red-shift by 75 nm in fluorescence spectrum upon binding to aggregated Aβ1-4_2_ fibrils as describes as previous reports [26]. Importantly, the probe exhibits little binding to other proteins involved with Aβ plaque formation such as lysozyme [59]. Though, we observed a percentage of binding to bovine serum albumin that is a plasma protein in the circulation and found in Aβ plaques. This is a common phenomenon of Aβ binding probes such as BTA-1, PIB, florbetaben, florbetapir etc [60–62]. Costaining of CRANAD-2 to brain sections from arc A mice with 6E10/OC antibodies of Aβ deposition clearly showed close resemblance in both vascular and parenchymal Aβ deposits.

Given the specificity of CRANAD-2 for Aβ deposits and the fact that the probe generates a detectable MSOT signal upon Aβ fibril binding, we explored the capability of *in vivo* MSOT cerebral Aβ mapping. Probes for MSOT imaging ideally have an absorbance and emission peak at NIR or even NIR II range to allow for sufficient penetration depth and separation from deoxygenated and oxygenated hemoglobin which are abundant in biological tissue but which have lower absorption in the NIR region of light [16]. This has been considered in the design of most Aβ probes, apart from NIAD-4 and Congo Red which have absorption peaks in the red part of the spectrum and which require invasive imaging approaches [18]. Moreover, some probes, e.g. BODIPY [20] derivatives and Congo red [27], do not pass across the blood-brain barrier and thus have to be injected intrathecally. CRANAD-2 has a suitable spectrum (excitation 640 nm, emission 685 nm) [26]. In previous reports it was shown that CRANAD-2 passes the blood-brain barrier [26, 27]. This has motivated us to inject the probe in arcAβ mice, which shows parenchymal plaque and cerebral amyloid angiopathy accumulation concomitantly with cognitive impairments from 6 months-of-age, where Aβ plaque load increases with age [39].

The optical absorption and fluorescence contrast provided by CRANAD-2 allowed for cross-validation by simultaneous real-time recording of hybrid 3D vMSOT and 2D epifluorescence data [50], useful to understand the relation between the fluorescence, absorbance and spectral dependence of the signals generated from probe binding to targets [63]. We observed uptake and wash-out of probe with MSOT and epifluorescence imaging, though the degree of changes in MSOT signal was lower. The differences may be explained by the fact that epifluorescence lacks the depth resolving capacity thus effectively adds up signals originating from different cranial compartments (skull and scalp over the head) to provide a surface-weighted signal [64], while tomographic MSOT imaging allows to map the probe uptake in the brain more accurately in 3D. Thus, epifluorescence imaging merely reflects wash-out of the probe from circulation, while MSOT closer represents the actual cerebral kinetics.

To harness the full potential of volumetric imaging, we performed serial MSOT imaging in arcAβ and non-transgenic littermates. Co-registration of MSOT data with a T_2_-weighted structural MRI dataset [37, 55] allowed to quantify regional probe uptake. We observed a clear difference in brain accumulation of CRANAD-2. In our study, the cortical signals detected by using epifluorescence imaging and vMSOT in arcAβ mice in this study was in accordance with the known Aβ distribution, which is highest in dorsal areas like the cortex and decreases ventrally [36, 39, 42], and the validation from immunohistochemistry. Several factors influence the image quality of MSOT, such as the modeling accuracy of light propagation in living tissues [65]. Further application of regional based fluence correction and non-negative correction in the reconstruction and unmixing steps will enable more accurate quantification [66, 67].

In conclusion, we demonstrated the suitability of CRANAD-2 for detecting Aβ deposits in a mouse model of AD amyloidosis with epifluorescence and MSOT imaging. The approach offers further longitudinal monitoring of therapeutics effect targeting at Aβ and for unveiling the disease mechanism in animal models.

## Availability of data and material

The datasets generated and/or analyzed during the current study are available in the repository DOI zenodo: 10.5281/zenodo.3672403.

## Author Contributions

RN, AV, PA and JK conceived and designed the study; RN, AV, XD, ZC performed the experiments; RN, AV, XD, ZC, PA analyzed the data; RN, AV, XD, ZC, PA, JK interpreted the results; RN, AV and JK wrote the initial paper draft; all coauthors contributed constructively to writing and editing the manuscript.

## Acknowledgement

The authors acknowledge support from Mr Michael Reiss, Ms Marie Rouault at Institute for Biomedical Engineering, ETH & University of Zurich; Mr. Daniel Schuppli at Institute for Regenerative Medicine, University of Zurich; Dr. Joachim Hehl of the Scientific Center for Optical and Electron Microscopy (ScopeM) of the Swiss Federal Institute of Technology ETH Zurich.

## Funding

JK received funding from the Swiss National Science Foundation (320030_179277), in the framework of ERA-NET NEURON (32NE30_173678/1), Vontobel foundation, Olga Mayenfisch Stiftung, and the Synapsis foundation. RN received funding from the University of Zurich Forschungskredit (Nr. FK-17-052), and Synapsis foundation career development award (2017 CDA-03), Helmut Horten Stiftung and Jubiläumsstiftung von Swiss Life. PA acknowledges financial support from the Synapsis foundation. Funding from the European Research Council (grant number ERC-CoG-2015-682379 to DR) is also acknowledged.

## Notes

### Competing Interest Statement

The authors have declared no competing interest.

